# Mechano-arrhythmogenicity is enhanced during late repolarisation in ischemia and driven by a TRPA1-, calcium-, and reactive oxygen species-dependent mechanism

**DOI:** 10.1101/2021.08.20.457138

**Authors:** Breanne A Cameron, T Alexander Quinn

## Abstract

**Background:** Cardiac dyskinesis in regional ischemia results in arrhythmias through mechanically-induced changes in electrophysiology (‘mechano-arrhythmogenicity’) that involve ischemic alterations in voltage-calcium (Ca^2+^) dynamics, creating a vulnerable period (VP) in late repolarisation.

**Objective:** To determine cellular mechanisms of mechano-arrhythmogenicity in ischemia and define the importance of the VP.

**Methods and Results:** Voltage-Ca^2+^ dynamics were simultaneously monitored in rabbit ventricular myocytes by dual-fluorescence imaging to assess the VP in control and simulated ischemia (SI). The VP was longer in SI than in control (146±7 *vs* 54±8ms; *p*<0.0001) and was reduced by blocking K_ATP_ channels with glibenclamide (109±6ms; *p*<0.0001). Cells were rapidly stretched (10-18% increase in sarcomere length over 110-170ms) with carbon fibres during diastole or the VP. Mechano-arrhythmogenicity, associated with stretch and release in the VP, was greater in SI than control (7 *vs* 1% of stretches induced arrhythmias; *p*<0.005) but was similar in diastole. Arrhythmias during the VP were more complex than in diastole (100 *vs* 69% had sustained activity; *p*<0.05). In the VP, incidence was reduced with glibenclamide (2%; *p*<0.05), by chelating intracellular Ca^2+^ (BAPTA; 2%; *p*<0.05), blocking mechano-sensitive TRPA1 (HC-030031; 1%; *p*<0.005), or by scavenging (NAC; 1%; *p*<0.005) or blocking reactive oxygen species (ROS) production (DPI; 2%; *p*<0.05). Ratiometric Ca^2+^ imaging revealed that SI increased diastolic Ca^2±^ (+9±1%, *p*<0.0001), which was not prevented by HC-030031 or NAC.

**Conclusion:** In ischemia, mechano-arrhythmogenicity is enhanced specifically during the VP and is mediated by ROS, TRPA1, and Ca^2+^.

## INTRODUCTION

Regional ischemia due to coronary artery occlusion is associated with deadly ventricular arrhythmias.^1^ Mechanical heterogeneity, acting through mechano-electric coupling mechanisms (‘mechano-arrhythmogenicity’), is thought to contribute to this arrhythmogenesis,^2^ and is supported by the strong correlation between regional ventricular wall motion abnormalities and arrhythmias in patients with coronary artery disease.^3^ In animal models, these arrhythmias have been shown to originate at the ischemic border,^4^ a site of systolic stretch of weakened ischemic myocardium.^5^ Consequently, arrhythmia incidence in ischemia is ventricular load-dependent,^4^ with distension of the ischemic region being a strong predictor of ventricular fibrillation.^6,7^ Computational modelling suggests that mechano-arrhythmogenicity in ischemia is the result of stretch-activated ion channel-mediated depolarisation at the ischemic border, which contributes to ectopic foci (if supra-threshold) or conduction slowing and block (if sub-threshold).^8^ Yet, the molecular identity of the mechano-sensitive ion channels involved remains unknown.^9^

Recent evidence from rabbit isolated heart studies suggests that ventricular mechano-arrhythmogenicity in regional ischemia is Ca^2+^-mediated,^10^ and may relate to a VP in late repolarisation during which a temporal dissociation between the recovery of membrane potential and cytosolic Ca^2+^ results in Ca^2+^ remaining elevated as myocytes become re-excitable.^10,11^ Further work showed that the mechano-sensitive,^12^ Ca^2+^-permeable^13^ transient receptor potential ankyrin 1 (TRPA1) channel^14^ can act as a source for Ca^2+^-mediated mechano-arrhythmogenicity in ventricular myocytes by triggering premature excitation and creating a substrate for more complex arrhythmic activity.^15^ Since the response of TRPA1 to mechanical stimulation is dependent on its baseline activity,^16^ which is increased in ischemia^17^ as it is agonised by multiple ischemic factors (*e.g*., increased cytosolic Ca^2+^ and ROS),^18,19^ TRPA1 may be involved in ischemic mechano-arrhythmogenicity.

Other mechano-sensitive processes may additionally contribute to arrhythmogenesis in ischemia. Stretch is known to increase NADPH oxidase-dependent ROS production (X-ROS)^20^ and subsequent sarcoplasmic Ca^2+^ release events *via* ryanodine receptors (RyR).^21^ As these mechanically-induced effects are enhanced in ischemia,^22^ regional stretch may lead to localised increases in ROS and cytosolic Ca^2+^, resulting in arrhythmic activity.^23^

The goal of this study was to investigate cellular mechanisms of ischemic mechano-arrhythmogenicity and the importance of the VP. Rabbit isolated ventricular myocytes exposed to simulated ischemic (SI) conditions were stretched in diastole and during the VP using a carbon-fibre-based system, combined with dual-parametric fluorescence imaging of voltage and cytosolic Ca^2+^, video-based measurement of sarcomere dynamics, and pharmacological interrogations. It was hypothesised that the incidence of stretch-induced arrhythmias would be greatest in the VP, and driven by a TRPA1-, Ca^2+^-, and ROS-mediated mechanism.

## METHODS

### Ethics

Experiments were conducted in accordance with the ethical guidelines of the Canadian Council on Animal Care with all protocols approved by the Dalhousie University Committee for Laboratory Animals. Details have been described following the Minimum Information about a Cardiac Electrophysiology Experiment (MICEE) reporting standard.^24^

### Ventricular myocyte isolation

Rabbit ventricular myocytes were enzymatically isolated as previously described.^15^ Details are available in the supplemental methods.

### Carbon fibre-based cell stretch

Cells were subjected to axial stretch using the carbon fibre (CF) method previously described for stretch of ventricular myocytes.^15,21^ Figure 2 shows a schematic for the stretch protocol. Contractile function and characteristics of stretch were assessed by monitoring sarcomere length and piezo-electric translators (PZT) and CF tip positions to determine incidence and classification of mechano-arrhythmogenicity (Fig. 2). Details can be found in the supplement.

**Figure 1.**
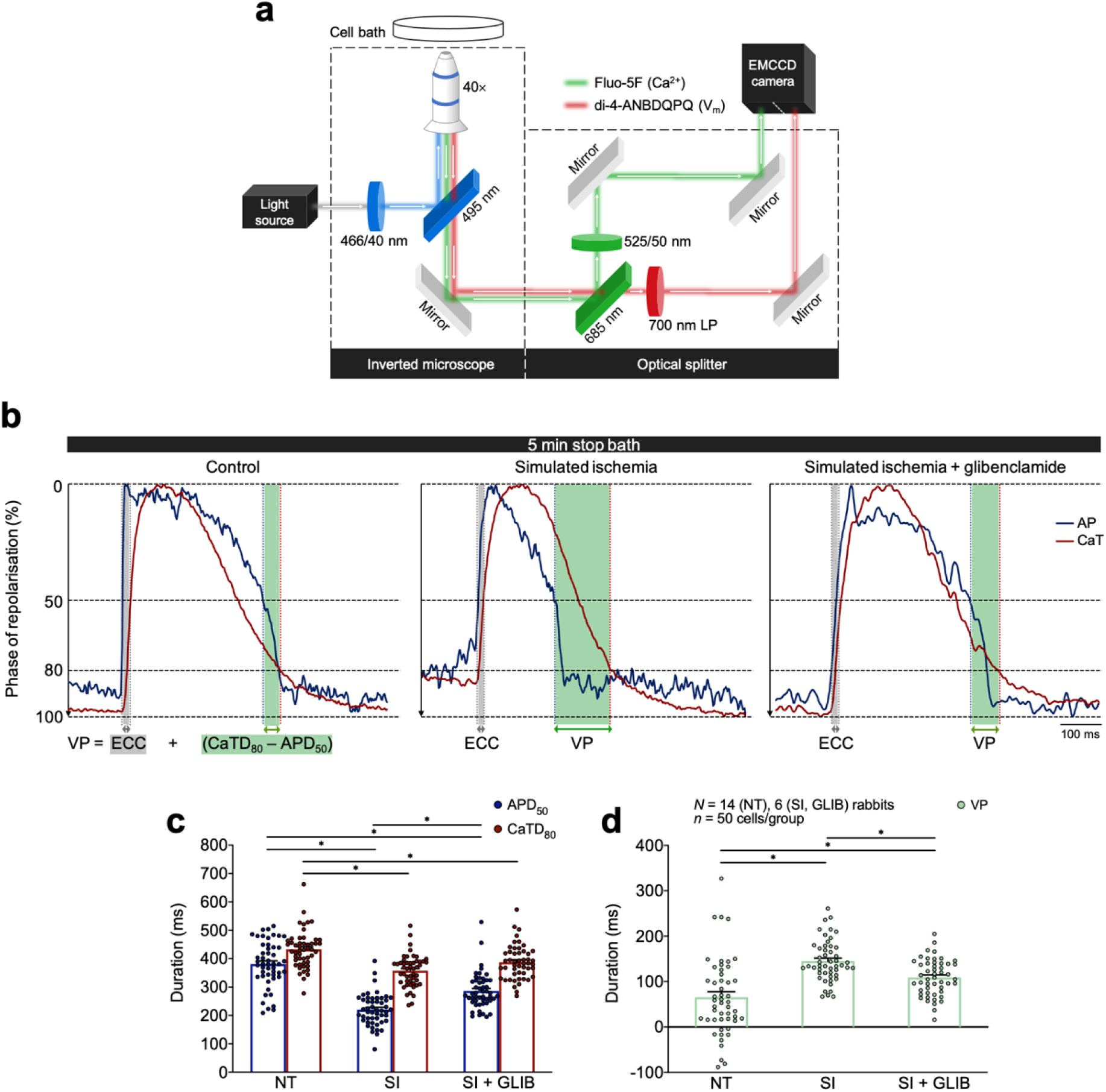
Temporal uncoupling of voltage-Ca^2+^ dynamics in ischemic ventricular myocytes is reduced by block of K_ATP_ channels with glibenclamide. **a**, Schematic of the single-excitation/dual-emission fluorescence imaging technique, utilising di-4-ANBDQPQ (20 μM for 14 min) and Fluo-5F-AM (5 μM for 20 min) indicators and a single camera-image splitter system. **b,** Trace of an action potential (AP, blue) and Ca^2+^ transient (CaT, red) simultaneously recorded in a contracting, paced (1 Hz) ventricular myocyte after 5 min exposure to either control (left), or to SI solution alone (middle) or following pre-incubation with glibenclamide (20 μM for 15 min, right). The calculated VP is shown in green. **c**, Average APD50 (blue) and CaTD80 (red) after 5 min in either normal Tyrode (NT), or in SI alone or with glibenclamide (SI + GLIB). **d**, Average VP in NT, SI, or SI + GLIB. Differences assessed by one-way ANOVA, with Tukey *post-hoc* tests. \**p*<0.05 between groups. Error bars represent SEM.

**Figure 2.**
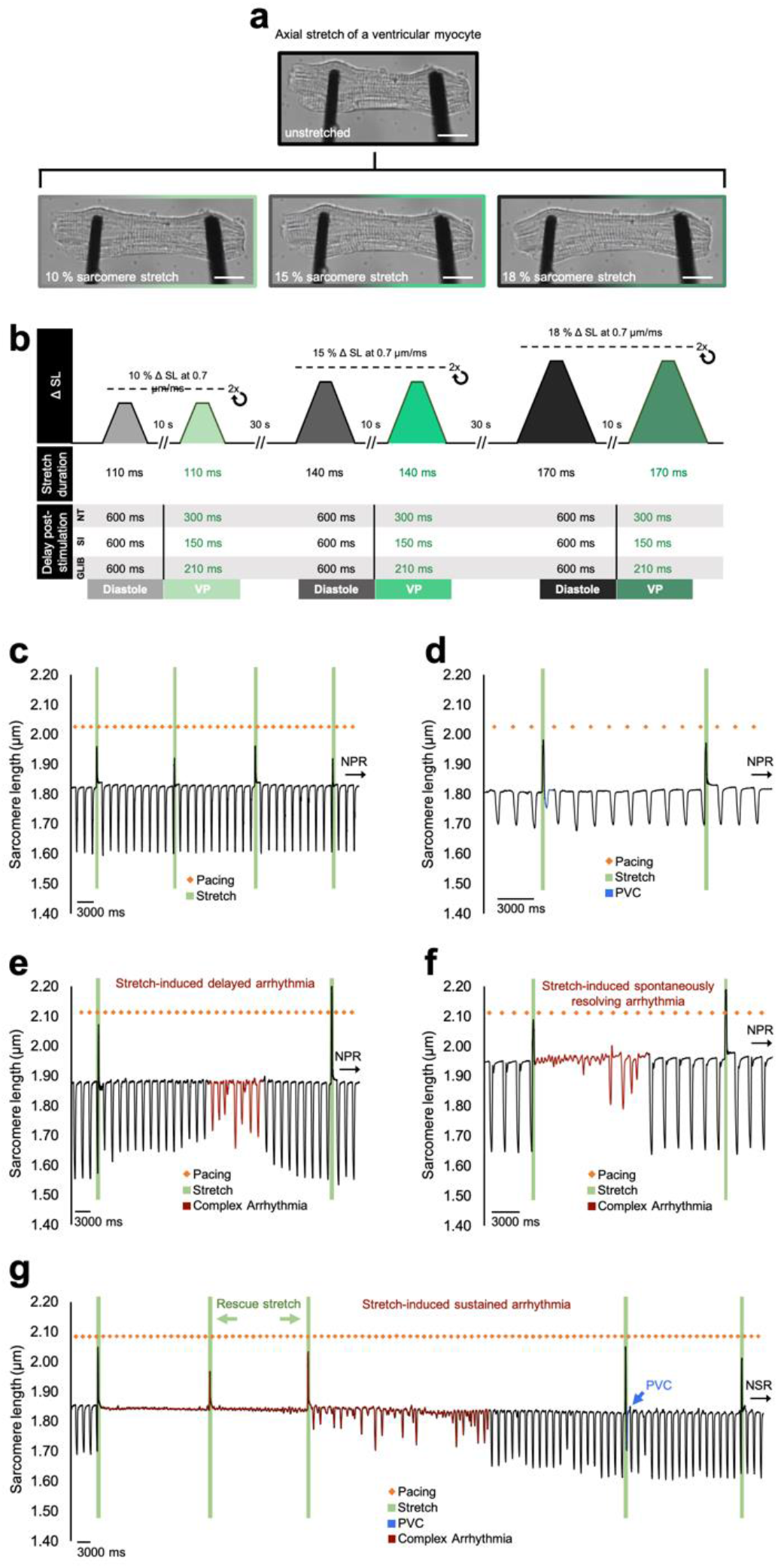
Protocol for timed transient stretch of ventricular myocytes and arrhythmia classification. **a**, Rabbit ventricular myocyte before (top) and during (bottom) unidirectional axial stretch using a carbon-fibre based system at increasing magnitudes of PZT displacement (left to right: 20, 30, and 40 μm). **b**, Schematic of the protocol for cell stretch timed in mid-diastole and the VP. **c**, Sarcomere trace of an ischemic cell (paced at 1 Hz, orange dots) that was stretched (green) and maintained normal paced rhythm (NPR). **d**, Stretch-induced premature ventricular contraction (PVC, blue segment). **e**, Stretch-induced delayed complex arrhythmia (red segment). **f**, Stretch-induced sustained arrhythmic activity that spontaneously resolved. **g**, Stretch-induced sustained arrhythmic activity that was terminated by rescue stretches.

### Pharmacology

Pharmacologic agents were dissolved in distilled water or dimethyl sulfoxide (DMSO) as appropriate. Agents included: BAPTA-AM (1 μM, 20 min pre-incubation; Abcam), dantrolene (1 μM, 5 min pre-incubation; Abcam), HC-030031 (10 μM, 30 min pre-incubation; Abcam), N-acetyl-L-cysteine (NAC; 10 mM, 20 min pre-incubation; Sigma), diphenyleneiodonium (DPI; 3 μM, 60 min pre-incubation; Abcam), or glibenclamide (20 μM, 15 min pre-incubation; Abcam).

### Fluorescence imaging

Figure 1 shows a schematic of the single-excitation/dual-emission fluorescence imaging technique to monitor voltage-Ca^2+^ dynamics in isolated ventricular myocytes, adapted from our previous work^15^. Ratiometric Ca^2+^ levels were assessed using Fura Red-AM (5 μM; AAT Bioquest). Detailed methodology is included in the supplement.

### Statistics

Statistics were performed using GraphPad Prism 9. Differences in arrhythmia incidence were assessed using chi-square contingency tables and Fisher’s exact test. Differences were assessed by two-tailed, paired or unpaired Student’s t-test (for normally distributed data) or Wilcoxon matched-pairs test (for data that was not normally distributed), one-way ANOVA with Tukey *post-hoc* tests (for normally distributed data), or Kruskal-Wallis with Dunn’s multiple comparisons test (for non-normally distributed data), where appropriate. A *p*-value of < 0.05 was considered significant. The relevant test and number of replicates is indicated in each figure caption (*N* = rabbits, *n* = cells, *m* = stretches, *c* = complex arrhythmias).

## RESULTS

### Simulated ischemia creates a VP that can be reduced by blocking K_ATP_ channels

The fluorescence imaging approach for APD and CaTD (Fig. 1a) revealed that exposure to SI (5 minutes) decreased both APD (APD_50,NT_ = 381±11 ms *vs* APD_50,SI_ = 221±8 ms; *p*<0.0001) and CaTD (CaTD_80,NT_ = 433±9 ms *vs* CaTD_80,SI_ = 358±8 ms; *p*<0.0001) compared to control (Fig. 1b, c and Supplemental Fig. 1a, b). This increased the VP (= (CaTD_80_ – APD_50_) + ECC) in ischemic cells (VP_NT_ = 66±12 ms *vs* VP_SI_ = 145±6 ms; *p*<0.0001; Fig. 1d). The SI-induced decrease in APD was attenuated by pre-incubation with the K_ATP_ antagonist glibenclamide (APD_50,GLIB_ = 282±8 ms; *p*<0.0001; Fig. 1b, c, Supplemental Fig. 1c), with no effect on CaTD (CaTD_80,GLIB_ = 384±8 ms), ultimately reducing the VP (VP_,GLIB_ = 109±6 ms; *p*<0.05; Fig. 1d).

### Transient stretch of ventricular myocytes results in premature contractions and complex arrhythmias

Stretch of ventricular myocytes with CFs (Fig. 2a, b) resulted in premature contractions (1-2 unstimulated contractions; Fig. 2d) and complex activity, including delayed transient rhythm disturbances (Fig. 2e) and sustained arrhythmic activity that either spontaneously resolved (Fig. 2f), or that was terminated by an additional stretch (Fig. 2g). To ensure arrhythmic activity was not the result of stretch-induced cellular damage, contractile function was measured before and after completion of a stretch-induced event, which showed no change, suggesting that cells were not damaged (Supplemental Fig. 2). Stretch characteristics were measured at increasing PZT displacements, corresponding with an increase in percent sarcomere stretch (10.7±1, 16±2, and 18.5±2%; *p*<0.005), stretched sarcomere length (2.04±0.02, 2.15±0.02, and 2.21±0.03 μm; *p*<0.0001), and stretch force (0.55±0.01, 0.80±0.05, and 0.98±0.05 μN; *p*<0.0001). Importantly, while stretch characteristics scaled with PZT displacement similarly in all cells, absolute values of stretch parameters varied between cells, reflecting physiological ventricular heterogeneity in intrinsic cell stiffness and response to an ischemic insult.^25^ Notably, the diastolic sarcomere length within a given cell was maintained after stretch at each PZT displacement, indicating that CF slippage and subsequent cell buckling did not occur (Supplemental Fig. 3). Surprisingly, increased PZT displacement did not correspond with an increase in arrhythmias (though a clear trend was present, Supplemental Fig. 4).

### Mechano-arrhythmogenicity is enhanced in late repolarisation in ischemic cells

To determine whether mechano-arrhythmogenicity is dependent on stretch timing, stretch was applied in mid-diastole or during the VP in control and ischemic cells. We showed that the incidence of mechano-arrhythmogenicity was increased in ischemic cells compared to control in the VP (6.8 *vs* 1.2% of stretches induced arrhythmias; *p*<0.005) but not in diastole. Additionally, in ischemic cells, arrhythmias in the VP were proportionally more complex than those generated in diastole (100 vs 69% of events had complex activity; *p*<0.05; Fig. 3). Fluorescence-based measurement of APD and CaTD during stretch was performed in a subset of cells to assess the temporal relation of the stretch pulse to the VP, and its effect on arrythmia incidence. This showed that 96% of cases had either stretch, stretch-release, or both occur within the cell-specific VP, with 4% of stretches missing it entirely (Supplemental Fig. 5a). Of those cases with some portion of the stretch pulse within the VP, 52% had both stretch and release within the VP, 41% only had release, and 7% of cells only stretch occurred in the VP (Supplemental Fig. 5b). Yet, 100% of cases that resulted in a mechanically-induced arrhythmia were attributed to both stretch and release within the VP (which represented 50% of those stretches; Supplemental Fig. 5b, c). We then sought to determine whether mechano-arrhythmogenicity in ischemic cells was affected by reduced VP duration by blocking K_ATP_ channels (Fig.1b, d). Pre-incubating ischemic cells with glibenclamide reduced arrythmia incidence in the VP compared to ischemia alone (2.1 *vs* 6.8% of stretches; *p*<0.05) with no effect on diastolic incidence (Fig. 3).

**Figure 3.**
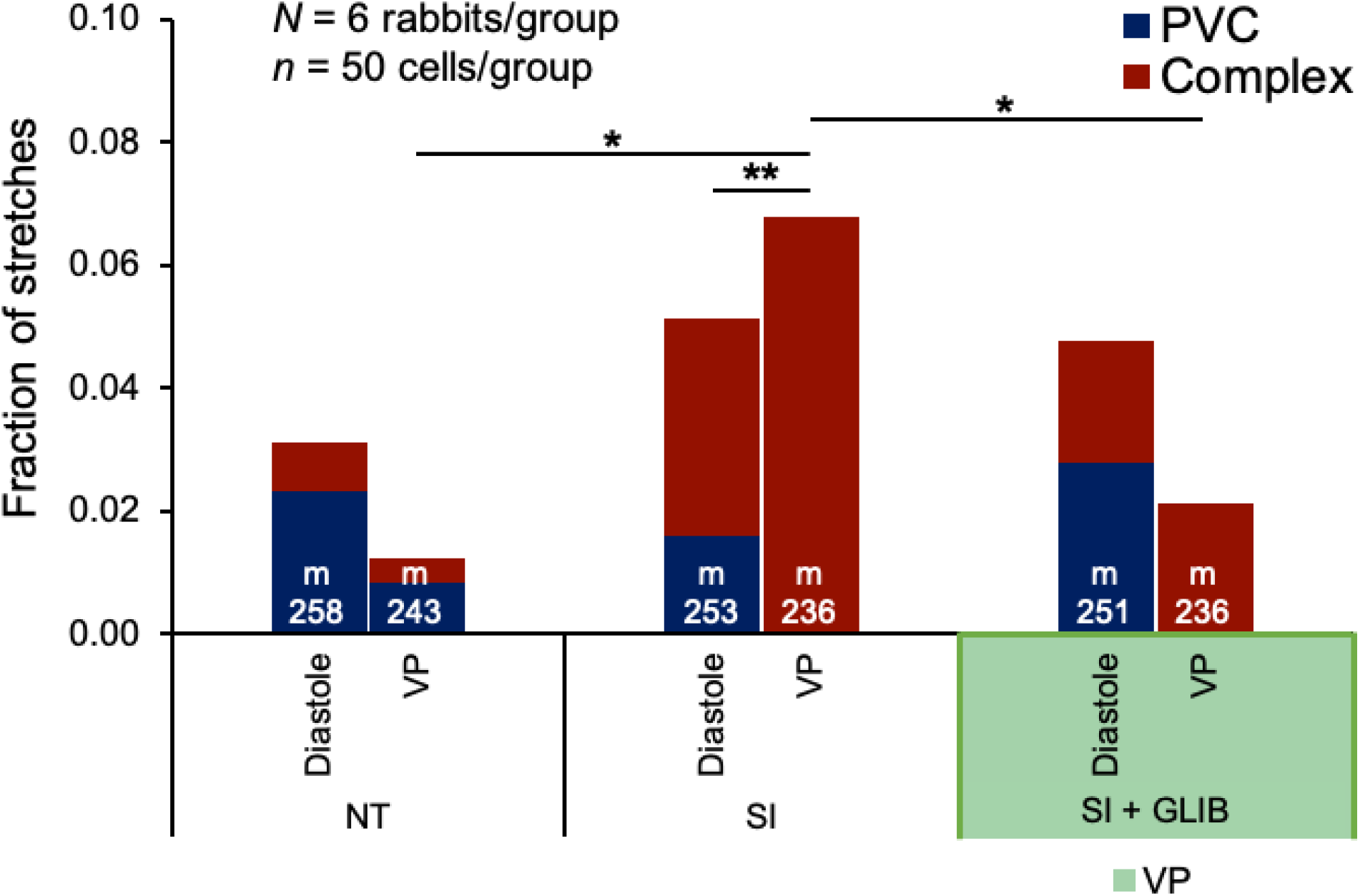
Role of the VP in ischemic mechano-arrhythmogenicity. Incidence of premature ventricular contractions (PVC, blue) and complex arrhythmias (red) with stretch during diastole or the VP after exposure to 5 min of normal Tyrode (NT), or to either SI alone or following pre-incubation with glibenclamide (20 μM for 15 min, SI + GLIB). Differences in arrhythmic incidence assessed using chi-square contingency tables and Fisher’s exact test. \**p*<0.05 between groups, \*\**p*<0.05 between diastolic and VP complexity within a group.

### TRPA1 channels and ROS mediate ischemic mechano-arrhythmogenicity during the VP

We next sought to determine mechanisms underlying the observed increase in mechano-arrhythmogenicity during the ischemic VP. As we previously showed that TRPA1 channels can act as a source for mechano-arrhythmogenicity,^15^ and as it is known that TRPA1 channel activity is increased in ischemia,^17^ we tested the effects of a specific TRPA1 blocker on arrythmia incidence. Pre-incubating ischemic cells with HC-030031 reduced arrhythmia incidence in the VP (0.9 *vs* 6.8%; *p*<0.005;), with no change in diastolic incidence (Fig. 4a), suggesting a role for TRPA1 channels in ischemic mechano-arrhythmogenicity.

**Figure 4.**
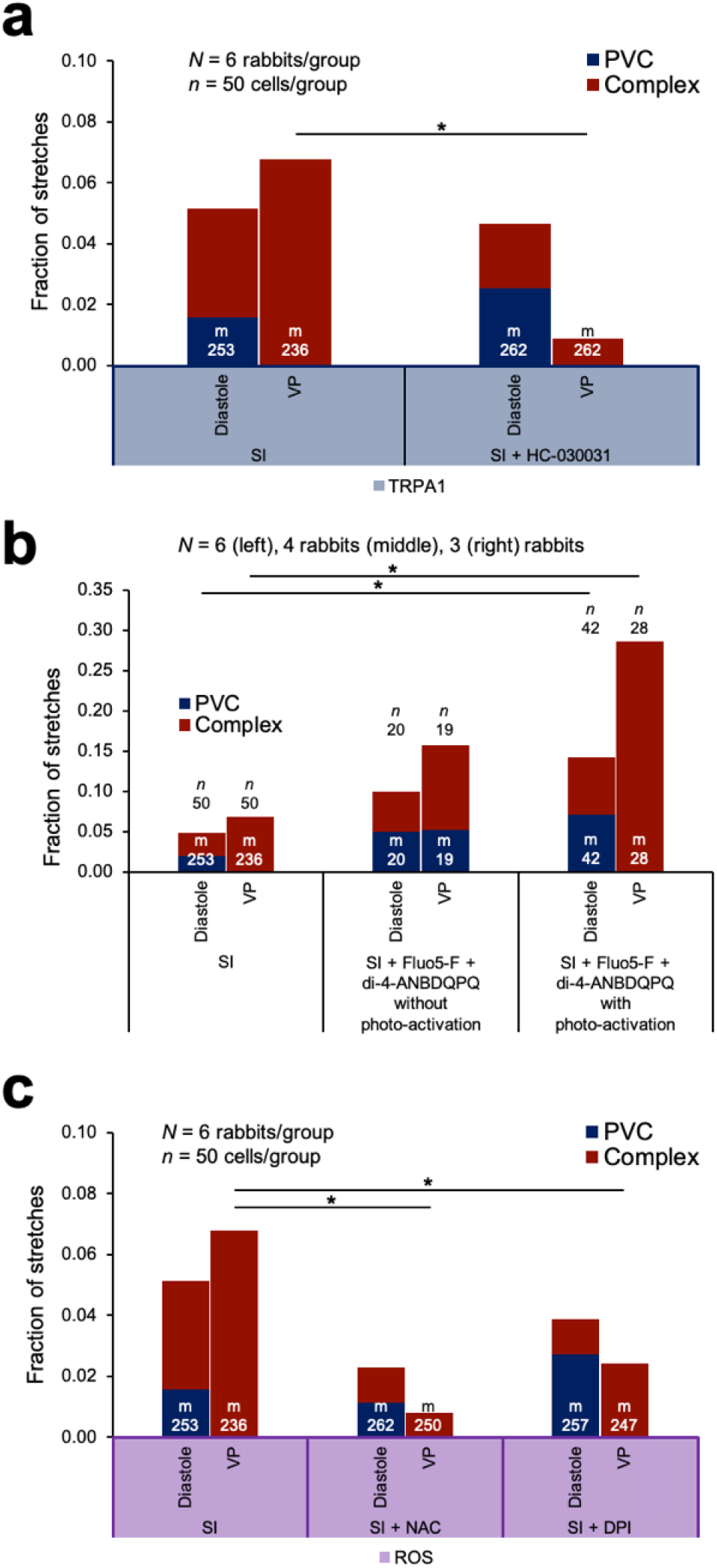
Role of ROS and TRPA1 channels in ischemic mechano-arrhythmogenicity. **a**, Incidence of premature ventricular contractions (PVC, blue) and complex arrhythmias (red) with stretch during diastole or the VP following pre-incubation with HC-030031 (10 μM for 30 min) and exposure to 5 min of SI. **b**, Incidence of arrhythmias with stretch during diastole or the VP in ischemic cells incubated with voltage (di-4-ANBDQPQ, 20 μM for 14 min) and Ca^2+^ (Fluo-5F-AM, 5 μM for 20 min) indicators, without (middle) or with (right) photo-activation. c, Incidence of arrhythmias during diastole or the VP in cells exposed to 5 min of SI pre-incubated with either NAC (10 mM for 20 min), or DPI (3 μM for 60 min). Differences in arrhythmic incidence assessed using chi-square contingency tables and Fisher’s exact test. \**p*<0.05 between groups.

Interestingly, while fluorescence imaging our ischemic cells, we observed a potentiated mechano-arrhythmogenicity during the VP (28.6 *vs* 6.8%; *p*<0.005), as well as an increase in diastole (14.3 *vs* 4.9%; *p*<0.05; Fig. 4b). However, this increase was only greater when the cells were exposed to photo-activation of the fluorescent dyes (Fig. 4b). As fluorophore photo-activation is associated with the generation of ROS,^26^ which is increased in ischemia,^22^ ROS might also play a role in mechano-arrhythmogenicity in ischemic cells. To test this, we pre-incubated ischemic cells with NAC to chelate intracellular ROS and found that arrhythmia incidence in the VP decreased compared to ischemia alone (0.8 *vs* 6.8%; *p*<0.0005), with no effect in diastole (Fig. 4c). Further, as we have shown that X-ROS production is enhanced in ischemia,^22^ we tested the effect of the NOX2 inhibitor DPI to block X-ROS production in ischemic cells. DPI also reduced arrhythmia incidence in the VP (2.4 vs 6.8%; *p*<0.05) with no diastolic effect (Fig. 4c). Combined, these data suggest that ROS, possibly through its effects on TRPA1,^19^ is a mediator of ischemic mechano-arrhythmogenicity.

### Cytosolic Ca^2+^ loading in ischemia is necessary, but not sufficient for enhanced mechano-arrhythmogenicity during the VP

As we have previously shown that increased cytosolic Ca^2+^ is necessary for TRPA1-mediated mechano-arrhythmogenicity in rabbit ventricular myocytes,^15^ we sought to assess whether it mediates the observed stretch-induced arrhythmias. We first buffered cytosolic Ca^2+^ in ischemic cells with BAPTA-AM and found that arrhythmia incidence was decreased in the VP (2.4 *vs* 6.8%; *p*<0.05; Fig. 5a). While this supports a necessary role for Ca^2+^ in ischemic mechano-arrhythmogenicity, surprisingly, arrhythmia incidence in these cells was simultaneously increased with stretch in diastole compared to control (8.5 *vs* 3.1%;*p*<0.05; Fig. 5a). Next, we assessed whether Ca^2+^ release *via* RyR may be involved in the observed arrhythmogenic role of cytosolic Ca^2+^ (like X-ROS, stretch-induced Ca^2+^-sparks are also enhanced by ischemia).^22^ To test this, RyR in ischemic cells were stabilised in their closed state with dantrolene, however, this had no effect on arrhythmia incidence with stretch in diastole or the VP (Fig. 5a). Finally, as Ca^2+^ loading in ischemia itself can be arrhythmogenic,^27^ we wanted to assess changes in cytosolic Ca^2+^ levels in our model. Through ratiometric Ca^2+^ imaging (using Fura Red-AM), we found that diastolic Ca^2+^ was increased with ischemia exposure (+9.3±1.1% change *p*<0.0001). Yet, while block of TRPA1 (with HC-030031) or chelation of ROS (with NAC) decreased mechano-arrhythmogenicity in ischemia, they did not prevent the increase in cytosolic Ca^2+^ (+10.0±1.0 or +12.3±1.4%), suggesting that neither TRPA1 nor ROS are solely responsible for the elevated Ca^2+^ levels in our ischemic cells (Fig. 5b).

**Figure 5.**
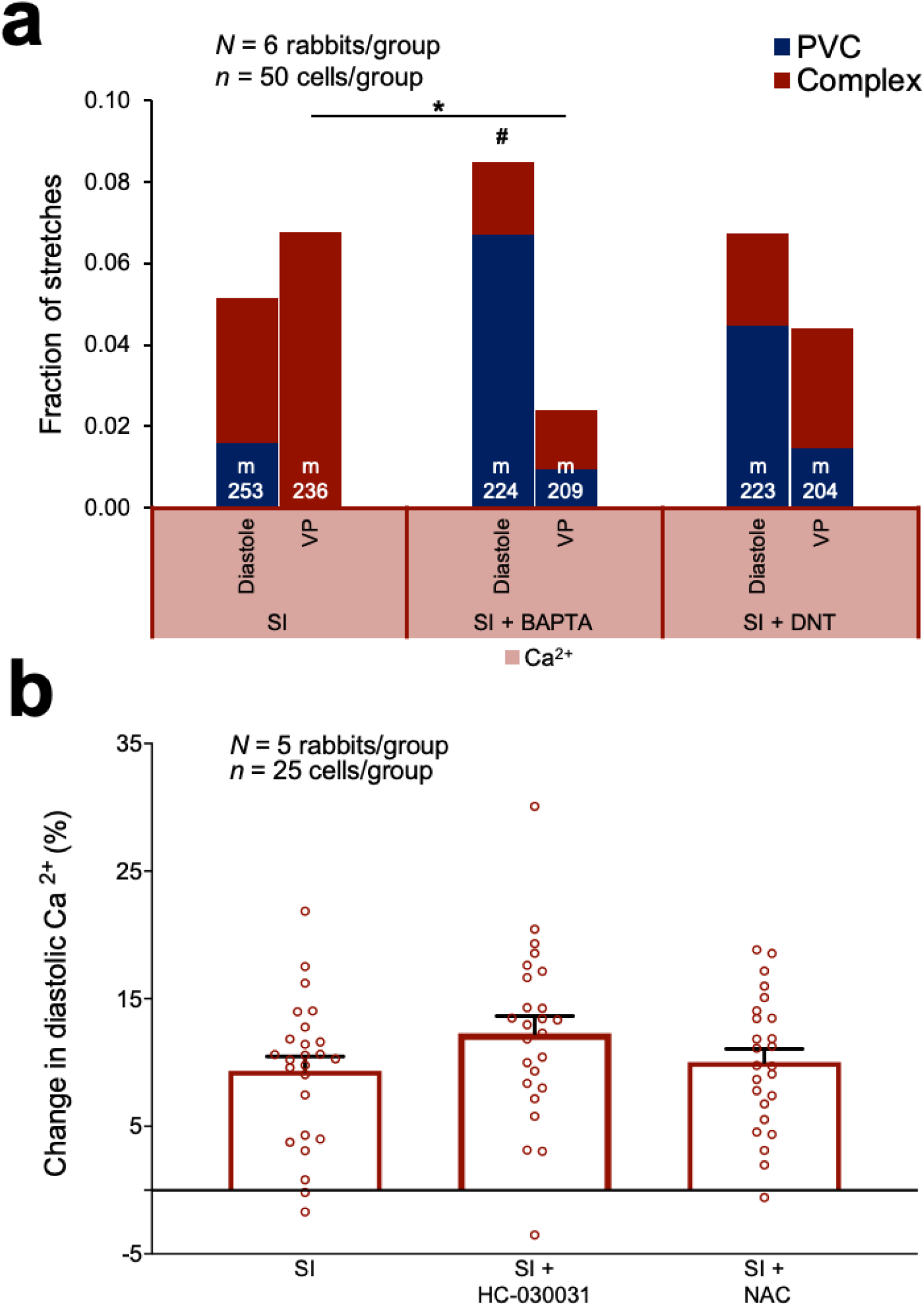
Role of intracellular Ca^2+^ in ischemic mechano-arrhythmogenicity. **a**, Incidence of premature ventricular contractions (PVC, blue) and complex arrhythmias (red) with transient stretch during diastole or the VP in cells exposed to 5 min of SI pre-incubated with BAPTA (middle; 1 μM for 20 min), or with dantrolene (right; DNT, 1 μM for 5 min). Differences in arrhythmic incidence assessed using chi-square contingency tables and Fisher’s exact test. \**p*<0.05 between groups, #*p*<0.05 compared to NT group. **b**, Percent change in diastolic Ca^2+^ from control after 5 min exposure to SI, SI + NAC (10 mM for 20 min pre-incubation), or SI + HC-030031 (10 μM for 30 min pre-incubation). Differences assessed using unpaired Student’s t-test between groups. Error bars represent SEM.

## DISCUSSION

In this study, we aimed to define cellular mechanisms of mechano-arrhythmogenicity and the importance of the VP in ischemia using rabbit ventricular myocytes exposed to SI and subjected to controlled stretch. We showed that mechano-arrhythmogenicity was enhanced only in the VP during SI, that arrhythmias generated in the VP were more complex than those in diastole, and that arrhythmogenesis involved TRPA1, cytosolic Ca^2+^, and ROS.

### Role of the VP in Ca^2+^-mediated mechano-arrhythmogenicity during ischemia

In the whole heart, ischemia-induced K_ATP_ activation causes a larger decrease in APD than CaTD, resulting in a VP for Ca^2+^-mediated arrhythmic activity.^11,27^ Elevated cytosolic Ca^2+^ can be arrhythmogenic by driving forward-mode sodium/Ca^2+^-exchanger (NCX) activity, approaching the threshold for premature excitation.^28^ Indeed, it has been shown that the generation of a VP through pharmacological K_ATP_ activation^11,15^ facilitates arrhythmogenesis. In ischemia, arrhythmogenicity of the VP may be exacerbated by transmural heterogeneity of K_ATP_ channel expression.^29^ Acute stretch may further increase cytosolic Ca^2+^ *via* Ca^2+^ influx through mechano-sensitive channels,^30^ which may also drive depolarisation and premature excitation by sodium influx.^9^ Furthermore, stretch has been shown to increase the affinity of myofilaments for Ca^2+^, such that systolic stretch causes excess myofilament Ca^2+^ loading.^30^ Upon release, dissociation of myofilament-bound Ca^2+^ can produce a surge in cytosolic Ca^2+^, which may induce SR Ca^2+^ release and generate Ca^2+^ waves, or drive NCX-mediated membrane depolarisation.^31^ Combined, these mechanisms may be sufficient to drive mechano-arrhythmogenicity in the VP.

In this study, we demonstrated the emergence of a VP in cells exposed to SI (Fig. 1). When transient stretch was timed to the VP, there was an increase in arrhythmia incidence that did not occur with stretch in diastole (Figure 3), revealing a temporal-dependence of ischemic mechano-arrhythmogenicity. Reducing the VP by blocking K_ATP_ channels (Fig. 1) resulted in an associated reduction in arrythmia incidence (Fig. 3), further supporting the role of the VP. The idea of whether stretch, stretch-release, or both, is critical to mechano-arrhythmogenicity remains an open question. However, our fluorescence measurements of APD, CaTD, and the VP during stretch revealed that all stretches that resulted in an arrhythmia stretched and released within the VP (Supplemental Fig. 5), suggesting that this combination may be necessary for mechano-arrhythmogenicity. This could partly explain why we did not see a higher incidence of arrhythmias in ischemic cells, as well as why reducing (but not eliminating) the VP attenuated arrhythmia incidence.

### Role of mechano-sensitive TRPA1 channels in ischemic mechano-arrhythmogenicity

A principal question underlying mechano-arrhythmogenicity is the identity of the mechano-sensitive ion channels involved.^9^ We demonstrated that TRPA1 channels can act as a source for Ca^2+^-mediated mechano-arrhythmogenicity.^15^ The potential role for TRPA1 channels in stretch-induced arrhythmias during ischemia is supported by their inherent mechano-sensitivity,^12^ preferential permeability to Ca^2+^,^13^ and their activation by ischemic factors, namely ROS^19^ and Ca^2+^,^18^ (which is bimodal, such that increased cytosolic Ca^2+^ enhances inward current to a point, after which greater increases in Ca^2+^ begin to inactivate channels).^32^ To test for their involvement in our cells, the specific TRPA1 blocker HC-030031 was used, which reduced arrhythmia incidence specifically in the VP (Fig. 4), supporting the role of TRPA1 channels in ischemic mechano-arrhythmogenicity.

The finding that mechano-arrhythmogenicity was exclusively increased during late repolarisation may also be related to a specific property of TRPA1 channels. While TRPA1 has been classically considered a non-voltage dependent channel, it has recently been shown that, under normal conditions, it is activated and inactivated by voltage at potentials outside the physiological range (+90 to +170 mV) so that voltage is not a relevant factor for its kinetics.^33^ However, when exposed to non-electrophilic agonists^34^ or elevated Ca^2+^,^18^ there is a leftward shift in its voltage activation into the physiological range. Further, continual TRPA1 agonism has been shown to de-sensitise the channel to Ca^2+^-mediated inhibition, effectively resulting in a sensitised channel with a physiological voltage dependence.^34^ Thus, in ischemia, increases in cytosolic Ca^2+^ and ROS may not only increase TRPA1 channel activity directly, but also indirectly through reduced Ca^2+^-mediated inhibition and modulation of its voltage dependence.

### Role of intracellular ROS in ischemic mechano-arrhythmogenicity

ROS production is increased in ischemia,^22^ which may contribute to mechano-arrhythmogenicity by increasing TRPA1 activity,^19^ and thus its response to mechanical stimulation.^16^ The potential mechanistic role of ROS in our ischemic cells was first revealed in our fluorescence imaging experiments, as fluorophore photo-activation, which generates ROS,^26^ resulted in an overall increase in arrythmia incidence (Fig. 4b). To more directly investigate the contribution of ROS to ischemic mechano-arrhythmogenicity, intracellular ROS was scavenged with NAC, or its NOX2 production was blocked with DPI, both of which resulted in a decrease in arrythmia incidence in the VP (Fig. 4c). The importance of ROS for mechano-arrhythmogenicity may extend beyond its potential effect on TRPA1 activity. X-ROS has also been shown to modulate RyR Ca^2+^ release,^22^ so it may additionally contribute to mechano-arrhythmogenicity through effects on cytosolic Ca^2+^.

### Role of cytosolic Ca^2+^ in ischemic mechano-arrhythmogenicity

Ischemic mechano-arrhythmogenicity in the VP appears to be mediated by cytosolic Ca^2+^, which is increased in ischemia.^27^ This was supported by the observation that chelating cytosolic Ca^2+^ with BAPTA reduced arrhythmia incidence in the VP (although, paradoxically it increased premature contractions in diastole, perhaps due to an increase in the driving force for Ca^2+^ influx with cytosolic Ca^2+^ buffering; Fig. 5a) This suggests that not only is Ca^2+^ involved in stretch-induced arrhythmias during ischemia, but also their stretch-timing dependence. Ischemic potentiation of stretch-induced RyR release has been suggested to be arrhythmogenic by contributing to cytosolic Ca^2+^ load and NCX-mediated membrane depolarisation.^22^ However, dantrolene (a RyR stabiliser) had no effect on arrhythmia incidence, suggesting that mechano-sensitive RyR release was not playing a critical role in arrhythmogenesis. Another potential mechanism of increased cytosolic Ca^2+^ is its direct activation of TRPA1 channels.^18^ Importantly, as cytosolic Ca^2+^ was not reduced by HC-030031 or NAC (Fig. 5b), despite a reduction in the incidence of stretch-induced arrythmias (Figs. 4a, b), it appears that cytosolic Ca^2+^ is necessary, but not sufficient for ischemic mechano-arrhythmogenicity.

## CONCLUSION

Ultimately, the observed ischemic mechano-arrhythmogenicity within the VP appears to relate to an increase in TRPA1 channel activity, driven by elevated intracellular ROS and cytosolic Ca^2+^ levels. Targeting TRPA1 channels in SI may help prevent electrical dysfunction and myocardial damage.^17^ The same may be true in other pathologies associated with changes in cardiac mechanics and TRPA1 modulating factors,^33^ such as ventricular pressure overload, in which TRPA1 has been shown to be involved in pathological changes^35^ and mechano-arrhythmogenicity is thought to occur,^5^ making TRPA1 channels a novel anti-arrhythmic target with exciting therapeutic potential.

## STATEMENTS

## Acknowledgements

Carbon fibres were a gift from Jean-Yves LeGuennec. We thank G. Iribe and K. Kaihara for technical assistance with cell stretch and the ischemic model design and I. Uzelac for assistance with electronic LED control.

## Author contributions

B.A.C. and T.A.Q. designed the study, interpreted the data, and wrote the manuscript; B.A.C. performed the experiments and analysed the data. All authors read and approved the manuscript.

## SUPPLEMENTAL METHODOLOGY

### Ventricular myocyte isolation

Single ventricular myocytes were enzymatically isolated from female New Zealand White rabbits (2.1 ± 0.2 kg, Charles River) euthanised by overdose through injection of pentobarbital (140 mg/kg) and heparin (1,500 units/kg, Sigma-Aldrich) into the marginal ear vein, followed by swift cardiac excision, aortic cannulation, and Langendorff perfusion (20 mL/min, 3-roller Watson-Marlow pump) with normal Tyrode (NT, 37 °C) solution (containing, in mM: 120 NaCl, 4.7 KCl, 1.0 MgCl_2_, 1.8 CaCl_2_, 10 glucose, 10 HEPES [Sigma-Aldrich], with pH adjusted to 7.40 ± 0.05 with NaOH and an osmolality of 300 ± 5 mOsm/L) bubbled with 100 % oxygen, for 10 min. The perfusate was then switched to a Ca^2+^-free solution (containing, in mM: 117 NaCl, 10 KCl, 1 MgCl_2_, 10 creatine, 20 taurine, 5 adenosine, 2 L-carnitine, 10 glucose, 10 HEPES [Sigma-Aldrich], with pH adjusted to 7.40 ± 0.05 with NaOH and an osmolality of 300 ± 5 mOsm/L) with the addition of 0.018 mM EGTA (Sigma-Aldrich), for 5 min. To begin enzymatic digestion, the perfusate was changed to the digestion solution (5 min at 20 mL/min, Gilson minipuls 3 pump) comprised of Ca^2+^-free solution with the addition of 200 U/mL Collagenase II (Worthington Biochemical Corporation), 0.06 mg/mL Protease XIV (from *Streptomyces griseus*, Sigma Aldrich), and 100 μM CaCl_2_, followed by a reduction in the perfusion rate to 15 mL/min until the heart became flaccid (~10-12 min). The left ventricular free wall was then removed and placed into 50 mL of stop solution, comprised of Ca^2+^-free solution with 0.5 % BSA (Sigma Aldrich) and 100 μM CaCl_2_. The ventricle was agitated and filtered through a 300 μm nylon mesh. The filtered tissue was resuspended in fresh stop solution, re-agitated, and re-filtered. The cell solution was split into 2 mL microcentrifuge tubes (VWR) and kept at room temperature (~22 °C, minimum 10 min). For experimentation, 1 mL of the supernatant was replaced with NT and the cells were left to equilibrate (10 min). The supernatant was then replaced with 100 % NT.

### Carbon fibre technique

Cells were subjected to unidirectional axial stretch using a pair of CFs (12-14 μm in diameter) affixed to glass capillaries (1.12 mm inner / 2 mm outer diameter, World Precision Instruments) with cyanoacrylate adhesive and fastened in microelectrode holders (MEH820, World Precision Instruments) that were coupled to triaxial water hydraulic micromanipulators (MHW-103, Narishige) and mounted on linear PZT (P-621.1CD, Physik Instrumente). The left and right CF were trimmed to 1.2 mm (compliant, translating fibre) and 0.6 mm (stiff, stationary fibre) in length, respectively.^15,22^ CF position was controlled by a piezo amplifier / servo controller (E-665.CR, Physik Instrumente) driven by a voltage signal generated from a DAQ device (USB-6361; National Instruments) and dictated by custom LabVIEW routines (National Instruments). CF stiffness was calibrated with a force transducer system (406A, Aurora Scientific). Force was measured for a given PZT displacement and fitted by linear regression to the formula: stiffness = force / PZT displacement. CF bending was calculated by monitoring PZT and CF tip positions (recorded at 240 Hz, Myocyte Contractility Recording System, IonOptix) and applying the formula (CF bend = change in CF tip distance - change in distance between PZTs). Stretch force was assessed from these values (force = CF stiffness x CF bend).

### Cellular stretch

A drop of the cell-containing solution was added to an imaging chamber (RC-27NE2, Warner Instruments) mounted on an inverted fluorescence microscope (IX-73, Olympus) with a 40× objective (UPLFLN40X, Olympus). The chamber contained 1 mL of either NT or a simulated ischemic solution to mimic 30 min of acute ischemia (containing, in mM: 140 NaCl, 15 KCl, 1.8 CaCl_2_, 1 MgCl_2_, 10 HEPES, 1 NaCN, and 20 2-deoxyglucose to block oxidative phosphorylation and anaerobic glycolysis; pH adjusted to 6.5 with NaOH), maintained at 35 °C by a temperature controller (TC-344C, Warner Instruments).^22,25^ The coverslip on the bottom of the chamber was coated with 20 μL of poly-2-hydroxyethyl methacrylate (poly-HEMA, Sigma-Aldrich) to prevent cellular adhesion. Once the cells had settled, bipolar electrical field stimulation (1 Hz, SIU-102, Warner Instruments) was commenced and contracting cardiomyocytes that were rod-shaped and clearly striated with intact membranes were chosen at random. Following cellular selection, pacing was halted, and the CFs were positioned at the lateral ends of the long axis of a cell and gently lowered onto the cell membrane using the triaxial hydraulic micromanipulators. Electrostatic adhesion of the cell to the CFs was confirmed by raising the cell off of the coverslip. Once cell attachment was established, electrical stimulation was recommenced, and cells contracted against the CFs for ~1-2 min to improve adhesion. After 5 min in the stop bath, unidirectional transient stretch (~112-173 ms total duration) using the compliant fibre was applied at a specific delay post-electrical stimulation, such that it occurred in two distinct time points of the electrical cycle: in mid-diastole or during the VP (as determined in initial experiments by simultaneous voltage-Ca^2+^ fluorescence-based imaging in each treatment, described below). This was repeated once for a total of four stretches, as follows: (i) mid-diastole (600 ms delay after an electrical stimulus in all treatments), followed by a 10 s pause, and (ii) during the VP (delay adjusted to 300, 150, or 210 ms in NT, SI, or SI with glibenclamide, respectively) in late repolarisation, followed by a 10 s pause. This protocol was repeated at increasing magnitudes of PZT displacement (20, 30, and 40 μm, with an average 10±1, 16±1, and 18±2% change in sarcomere length, respectively), with 30 s between each increase in magnitude, for a total of 12 stretches (Fig. 2a, b).

### Assessment of mechano-arrhythmogenicity

Contractile function (diastolic sarcomere length, rate and percentage of sarcomere shortening) and characteristics of stretch (percent change in sarcomere length, stretched sarcomere length, and applied force) were assessed by monitoring sarcomere length and PZT and CF tip positions (as described above). Arrhythmic activity with stretch was classified from sarcomere measurements into either (i) premature ventricular contractions (PVC, 1 or 2 unstimulated contractions), or (ii) complex activity (including delayed arrhythmic activity and sustained arrhythmic activity that either spontaneously resolved or was terminated by an additional stretch; Fig. 2). When an arrhythmia occurred, the next stretch was delayed by the appropriate amount (either 10 or 30 s). To control for cellular damage, any stretch that resulted in CF slippage, or a sustained arrhythmia that could not be terminated by a maximum of 2 stretches was excluded.

### Dual parametric voltage-Ca^2+^ fluorescence imaging

The Ca^2+^-sensitive dye Fluo-5F, AM (5 μM, ThermoFisher Scientific) and Pluronic F-127 (0.02 %, dissolved in DMSO, Biotium) were added to a microcentrifuge tube containing cells in 50% NT (20 min). The supernatant was then replaced with fresh full Ca^2+^ NT, and the voltage-sensitive dye di-4-ANBDQPQ (20 μM, University of Connecticut Health Centre) dissolved in ethanol was added to the tube and incubated for 14 min. The supernatant was again replaced with fresh NT, probenecid (1 mM, Sigma-Aldrich) was added, and the cells were maintained in the dark at room temperature until use (maximum 1 hour). When ready for imaging, the solution containing dye-loaded cells was gently agitated with a transfer pipette and a small drop was added to 1 mL of the relevant solution (NT or SI) in the imaging chamber. CFs were adhered (as described above) to allow for proper positioning of cells, and to reduce motion with cellular contraction in the direction perpendicular to the imaging plane, allowing for imaging without the use of an excitation-contraction uncoupler. Fluorescence was excited by a mercury lamp (U-HGLGPS, Olympus) passed through a 466/40 nm bandpass filter (FF01-466/40, Semrock) and reflected onto the sample by a 495 nm dichroic mirror (FF495-Di03, Semrock). For simultaneous measurement of transmembrane voltage and intracellular Ca^2+^, each fluorescent signal was projected onto one-half of a 128 × 128-pixel, 16-bit electron-multiplying charge-coupled device (EMCCD) camera sensor (iXon3, Andor Technology) using an emission image splitter (Optosplit II; Cairn Research) and recorded at 500 fps with 2 ms exposure and maximum electron-multiplying gain. The two signals were split with a 685 nm dichroic mirror (FF685-Di02, Semrock) and Fluo-5F emission was collected with a 525/50 nm bandpass filter (FF03-525/50, Semrock) and di-4-ANBDQPQ emission with a 700 nm long-pass filter (HQ700lp; Chroma Technology). A schematic of the imaging setup is provided in Fig. 1a.

Analysis of voltage-Ca^2+^ signals was performed using custom Matlab routines (R2018a, MathWorks). Whole-cell fluorescence was averaged, a temporal filter (50 Hz low-pass Butterworth) was applied, and bleaching was eliminated by fitting diastolic fluorescence over time with a second-order polynomial function and subtracting the result. From these signals, time to 20, 30, 50, or 80 % recovery of the action potential (action potential duration, APD) or the Ca^2+^ transient (Ca^2+^ transient duration, CaTD) were averaged over 3 consecutive cardiac cycles (Supplemental Fig. 1). The VP was calculated as the difference between CaTD80 and APD50, a period during which myocytes start to become re-excitable while cytosolic Ca^2+^ remains elevated, plus the difference between the timing of the action potential and Ca^2+^ transient upstrokes (excitation-contraction coupling time, ECC): VP = (CaTD_80_ - APD_50_) + ECC (Fig. 1b, d).

For assessment of the mechanical phase around the VP during fluorescence imaging (Supplemental Fig. 5), cells loaded with the voltage and Ca^2+^ indicators were stretched twice during photoexcitation (40 μm PZT displacement) with a 10 s pause in-between: once in diastole, and once in the VP. APD, CaTD, and VP values in these cells were calculated as above, and cross-compared with the timing of stretch, release, both, or neither within the VP.

### Ratiometric Ca^2+^ fluorescence imaging

Ratiometric Ca^2+^ levels were assessed using the Ca^2+^ indicator Fura Red-AM (5 μM; AAT Bioquest). Cells loaded with the dye were incubated with Pluronic F-127 (0.02 %) and probenecid (1 mM, dissolved in DMSO) for 20 min. Excitation was induced using alternating pulses from two white light-emitting diodes (CFT-90-W; Luminus Devices) each with a bandpass filter (420/10 nm, FF01-420/10, Semrock; or 531/22 nm, FF02-531/22, Semrock) that were combined into the microscope excitation light path with a 455 nm dichroic mirror (AT455dc, Chroma Technology) and reflected onto the sample by a 562 nm dichroic mirror (T562lpxr, Chroma Technology). Fluorescence emission was measured through a 632/60 nm bandpass filter (AT635/60m, Chroma Technology) with an EMCCD camera at a rate of 500 fps, with 2 ms exposure time and maximum electron-multiplying gain. Light pulses and camera frame acquisition were synchronised with a custom control box (supplied by Dr. Ilija Uzelac, Georgia Institute of Technology) so that alternating frames corresponded to the signal generated by the two excitation wavelengths.

Analysis of intracellular Ca^2+^ was performed using custom Matlab routines. Whole-cell fluorescence was averaged, and a temporal filter (50 Hz low-pass Butterworth) was applied. The two Ca^2+^ signals were separated, and the ratio was calculated. Any remaining baseline drift was eliminated by fitting the resulting diastolic fluorescence signal with a second-order polynomial function and subtracting the result. From the corrected signals, the minimum value for each cardiac cycle (representing the diastolic Ca^2+^ level) was averaged over 3 consecutive cardiac cycles. To assess changes in intracellular Ca^2+^ following exposure to SI, cells were first imaged in NT to get a baseline value, followed by a change to SI solution by perfusion at 2.1 mL/min through an inline heater (SF-28, Warner Instruments) for 2 min. Perfusion was stopped, and cells were maintained in the ischemia solution for 5 min before the second measurements were recorded.

### Code availability

All custom computer source code used in this study is available from the corresponding author upon reasonable request.

### Data availability

The datasets generated during and/or analysed during the current study are available from the corresponding author upon reasonable request.

## SUPPLEMENTAL FIGURES

**Supplemental Figure 1.**
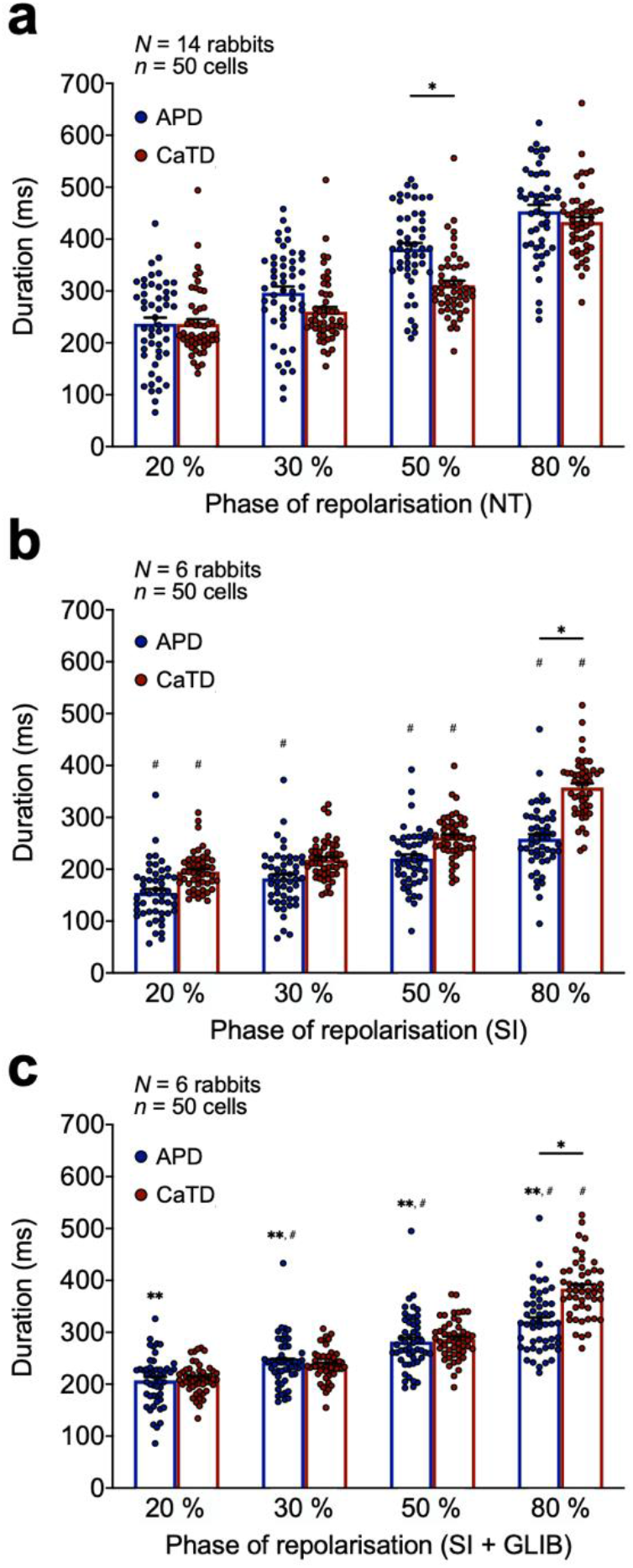
Effect of normal Tyrode or simulated ischemia alone or following pre-incubation with glibenclamide on cellular voltage-Ca^2+^ dynamics. **a,** Average APD (blue) and CaTD (red) at 20, 30, 50, and 80% repolarisation after 5 min in normal Tyrode (NT) measured using a fluorescence imaging technique with voltage (di-4-ANBDQPQ, 20 μM for 14 min) and Ca^2+^ (Fluo-5F-AM, 5 μM for 20 min) fluorescent indicators and a single camera-image splitter system. **b,** Average APD (blue) and CaTD (red) at 20, 30, 50, and 80 % repolarisation after 5 min in SI. **c,** Average APD (blue) and CaTD (red) at 20, 30, 50, and 80 % repolarisation after 5 min in SI following pre-incubation with glibenclamide (20 μM for 15 min, SI + GLIB). Differences assessed by one-way ANOVA, with Tukey *post-hoc* tests. \**p*<0.05 between groups, #*p*<0.05 compared to NT group, \*\**p*<0.05 compared to SI group. Error bars represent standard error of the mean.

**Supplemental Figure 2.**
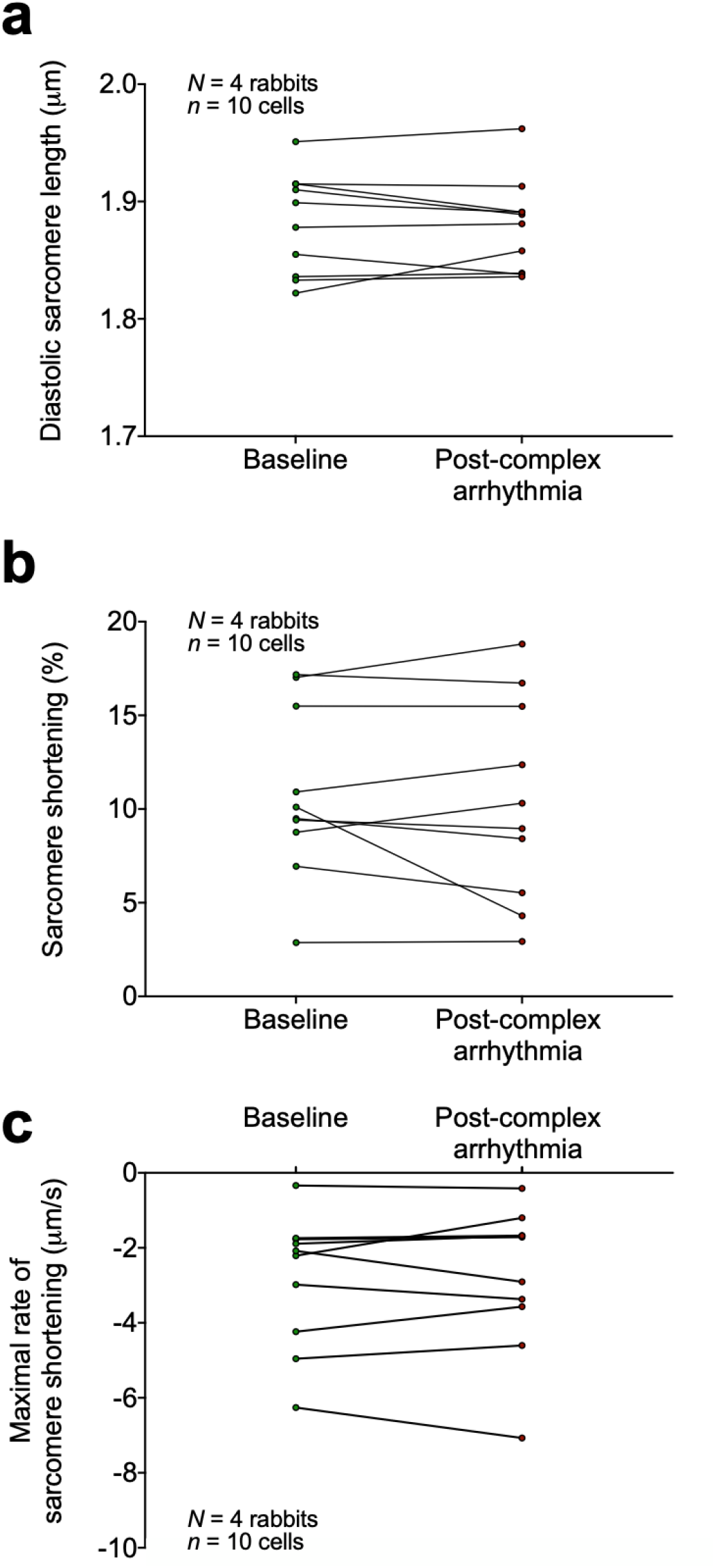
Effect of complex arrhythmias on contractile function. **a,** Diastolic sarcomere length **b,** percentage sarcomere shortening, and **c,** maximal rate of sarcomere shortening before and after resolution of a complex arrhythmia in ventricular myocytes exposed to 5 min of SI. Differences assessed by paired Student’s t-test.

**Supplemental Figure 3.**
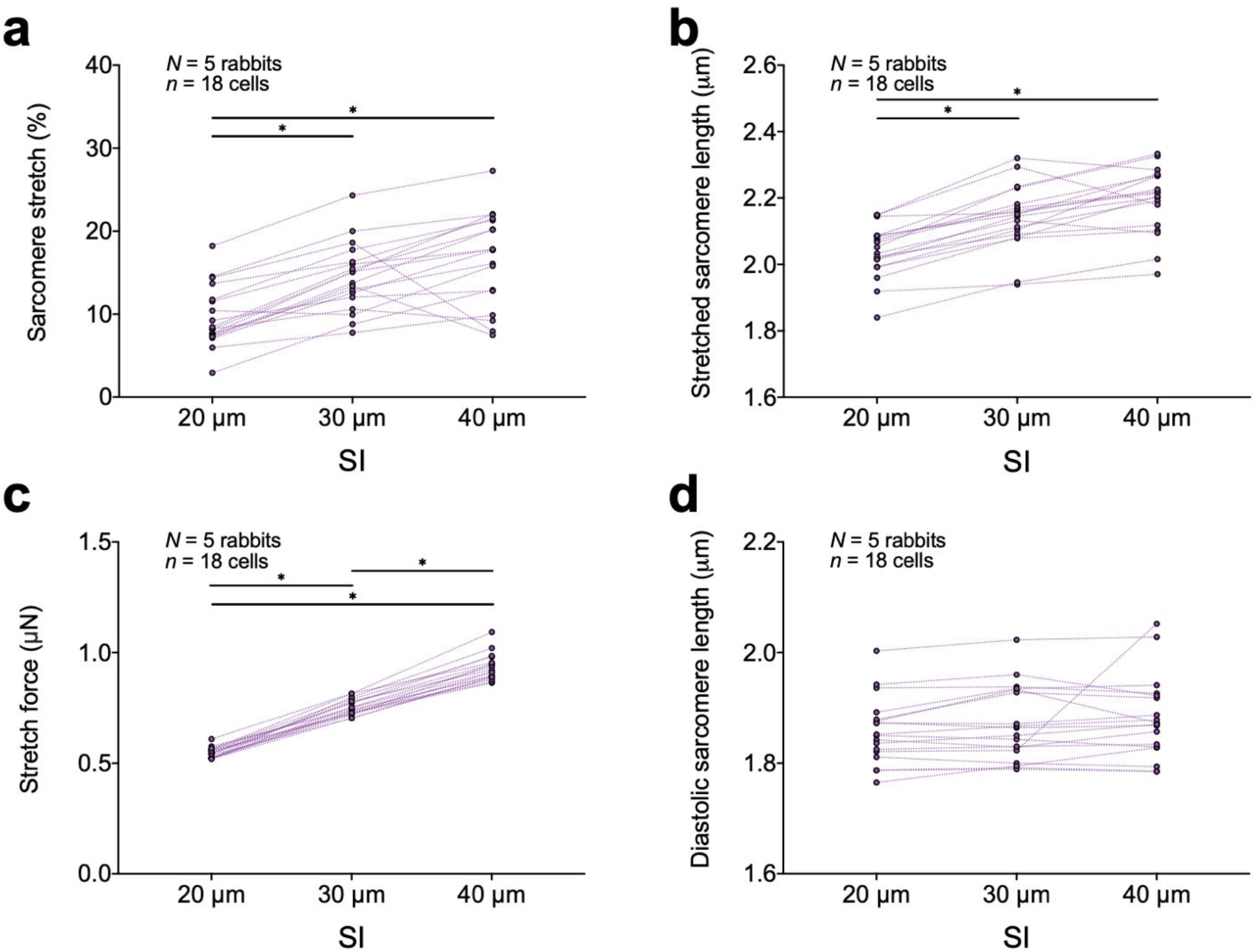
Mechanical parameters of cell stretch in ischemic cells. **a,** Percentage sarcomere stretch, **b,** maximal stretched sarcomere length **c,** maximal applied force during-, and **d,** diastolic sarcomere length following rapid, transient stretch of rabbit isolated ventricular myocytes exposed to 5 min of SI with increasing levels of PZT movement (20, 30, and 40 μm). Differences assessed by one-way ANOVA with Tukey *post-hoc* tests. \**p*< 0.05 within groups. Error bars represent standard error of the mean.

**Supplemental Figure 4.**
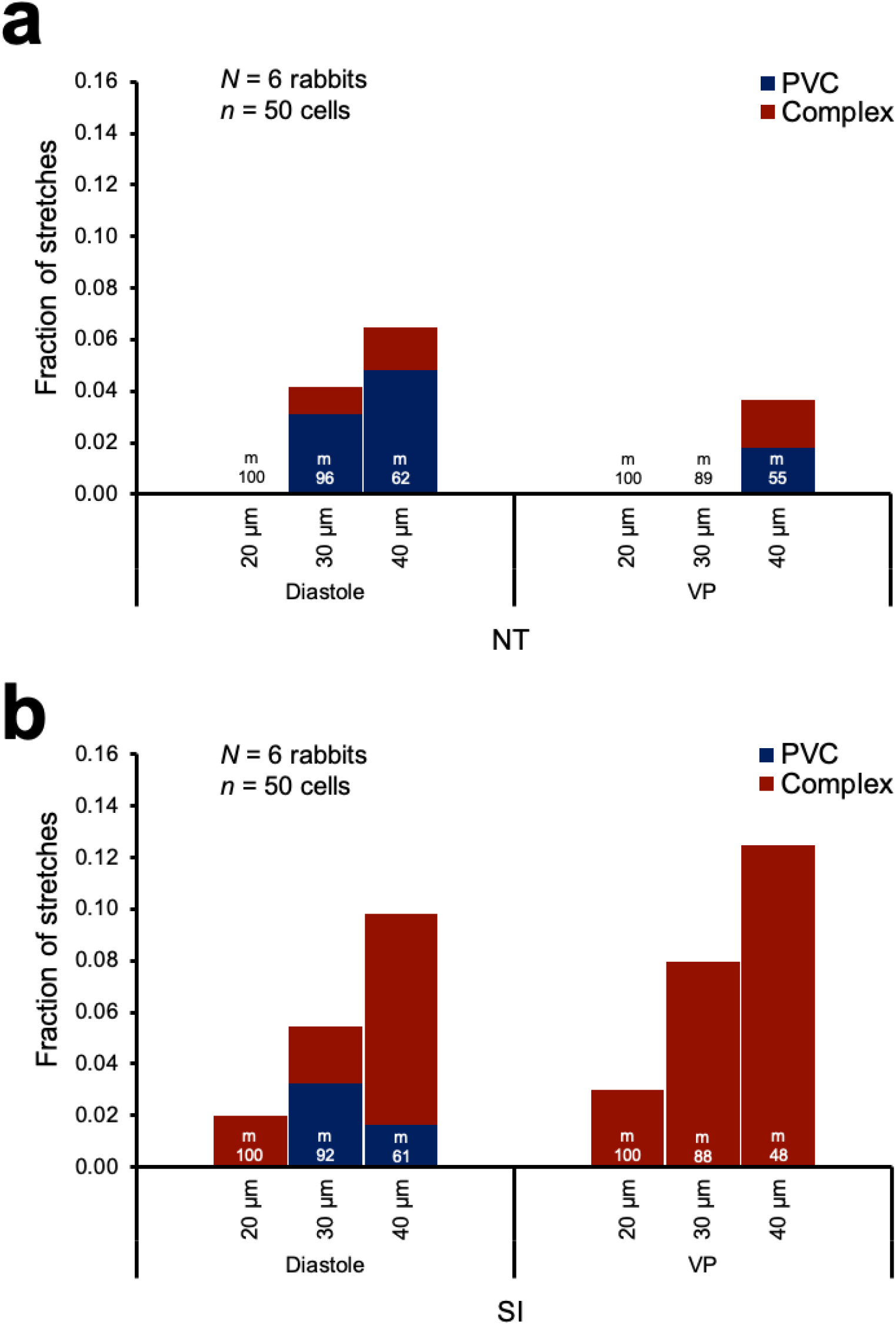
Effect of stretch magnitude on the incidence of mechano-arrhythmogenicity. **a,** Incidence of premature ventricular contractions (PVC, blue) and complex arrhythmias (red) with rapid, transient stretch during diastole (left) and the VP (right) with increasing levels of PZT movement (20, 30, and 40 μm) in rabbit isolated ventricular myocytes exposed to 5 min of normal Tyrode (NT). **b,** Incidence of arrhythmias in cells exposed to 5 min of SI. Differences assessed using chi-square contingency tables and Fisher’s exact test. *m* = stretches.

**Supplemental Figure 5.**
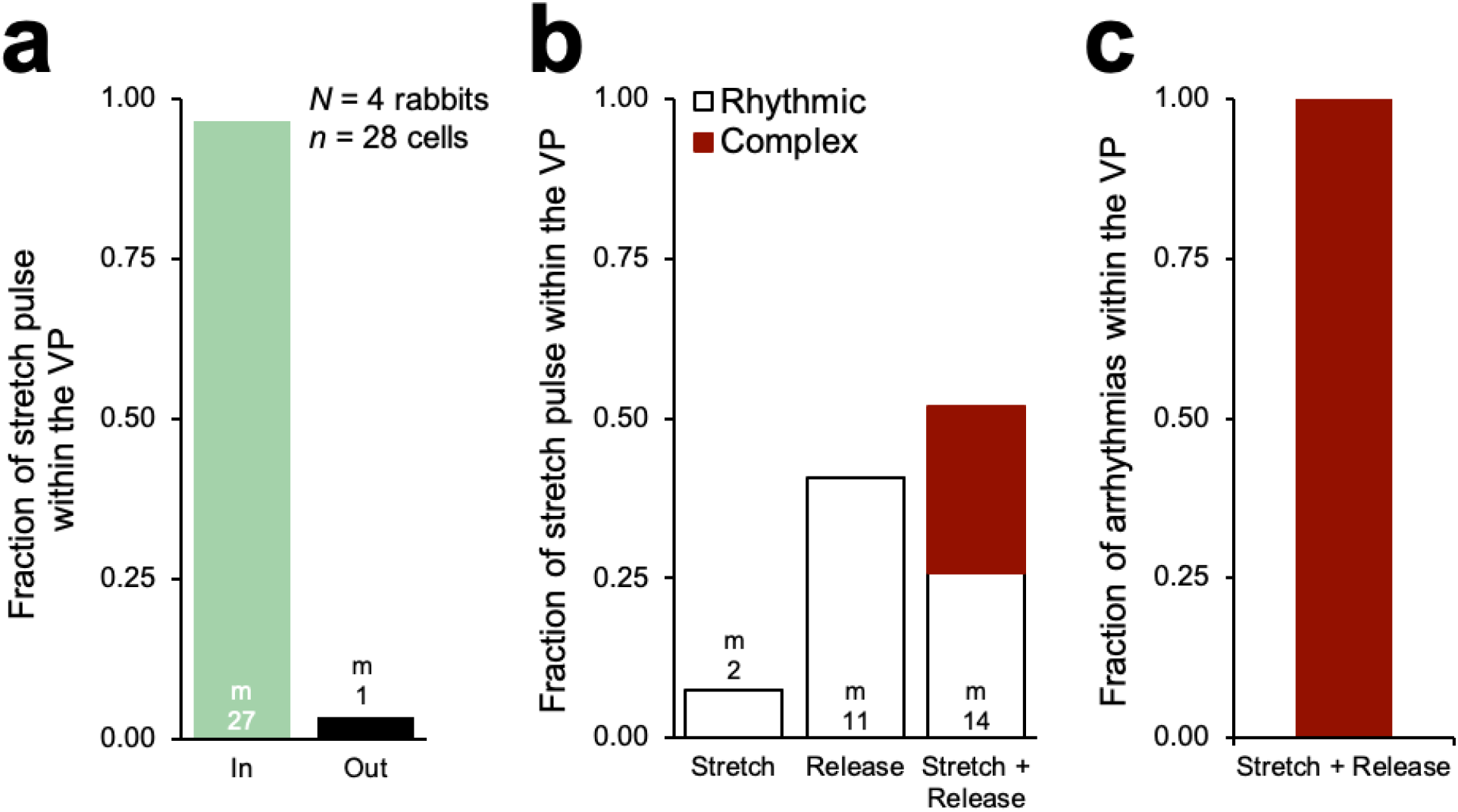
Role of stretch and/or release during the VP in ischemic mechano-arrhythmogenicity. **a,** Fraction stretch pulse segment (only stretch, only release, or both stretch and release) that occurred within (IN) or outside (OUT) the VP in ischemic cells, revealed by fluorescence imaging (di-4-ANBDQPQ, 20 μM for 14 min; and Fluo-5F-AM, 5 μM for 20 min) combined with timed stretch. **b**, Of the stretch pulse segments revealed to be within the VP, the fractions that were (i) stretch only, (ii) release only, or (iii) both. Associated arrhythmias shown in red. **d**, Fraction of resultant arrhythmias during the VP associated with (i) stretch only, (ii) release only, or (iii) stretch and release. *m* = stretches, *c* = complex arrhythmias.

## REFERENCES

1. Rubart M, Zipes DP. Mechanisms of sudden cardiac death. J Clin Invest 2005;115:2305–2315. doi:10.1172/jci26381.

2. Quinn TA, Kohl P. Cardiac mechano-electric coupling: acute effects of mechanical stimulation on heart rate and rhythm. Physiol Rev 2021;101:37–92. doi:10.1152/physrev.00036.2019.

3. Siogas K, Pappas S, Graekas G, Goudevenos J, Liapi G, Sideris DA. Segmental wall motion abnormalities alter vulnerability to ventricular ectopic beats associated with acute increases in aortic pressure in patients with underlying coronary artery disease. Heart 1998;79:268. doi:10.1136/hrt.79.3.268.

4. Coronel R, Wilms-Schopman FJG, deGroot JR. Origin of ischemia-induced phase 1b ventricular arrhythmias in pig hearts. J Am Coll Cardiol 2002;39:166–176. doi:10.1016/s0735-1097(01)01686-2.

5. Quinn TA. The importance of non-uniformities in mechano-electric coupling for ventricular arrhythmias. J Interv Card Electr 2014;39:25–35. doi:10.1007/s10840-013-9852-0.

6. Barrabés JA, Inserte J, Rodríguez-Sinovas A, Ruiz-Meana M, Garcia-Dorado D. Early regional wall distension is strongly associated with vulnerability to ventricular fibrillation but not arrhythmia triggers following coronary occlusion in vivo. Prog Biophysics Mol Biology 2017;130:387–393. doi:10.1016/j.pbiomolbio.2017.05.012.

7. Barrabés JA, Garcia-Dorado D, Padilla F, Agulló L, Trobo L, Carballo J, et al. Ventricular fibrillation during acute coronary occlusion is related to the dilation of the ischemic region. Basic Res Cardiol 2002;97:445–451. doi:10.1007/s003950200051.

8. Rice JJ, Winslow RL, Dekanski J, McVeigh E. Model studies of the role of mechano-sensitive currents in the generation of cardiac arrhythmias. J Theor Biol 1998;190:295–312. doi:10.1006/jtbi.1997.0538.

9. Peyronnet R, Nerbonne JM, Kohl P. Cardiac mechano-gated ion channels and arrhythmias. Circ Res 2016;118:311–329. doi:10.1161/circresaha.115.305043.

10. Baumeister PA, Lawen T, Rafferty SA, Taeb B, Uzelac I, Fenton FH, et al. Mechanically-induced ventricular arrhythmias during acute regional ischemia. J Mol Cell Cardiol 2018;124:87–88. doi:10.1016/j.yjmcc.2018.07.021.

11. Tang L, Joung B, Ogawa M, Chen P-S, Lin S-F. Intracellular calcium dynamics, shortened action potential duration, and late-phase 3 early afterdepolarization in Langendorff-perfused rabbit ventricles. J Cardiovasc Electr 2012;23:1364–1371. doi:10.1111/j.1540-8167.2012.02400.x.

12. Moparthi L, Zygmunt PM. Human TRPA1 is an inherently mechanosensitive bilayer-gated ion channel. Cell Calcium 2020;91:102255. doi:10.1016/j.ceca.2020.102255.

13. Bobkov YV, Corey EA, Ache BW. The pore properties of human nociceptor channel TRPA1 evaluated in single channel recordings. Biochim Biophys Acta 2010;1808:1120–1128. doi:10.1016/j.bbamem.2010.12.024.

14. Wang Z, Ye D, Ye J, Wang M, Liu J, Jiang H, et al. The TRPA1 channel in the cardiovascular system: promising features and challenges. Front Pharmacol 2019;10:1253. doi:10.3389/fphar.2019.01253.

15. Cameron BA, Stoyek MR, Bak JJ, Quinn TA. TRPA1 channels are a source of calcium-driven cardiac mechano-arrhythmogenicity n.d. doi:10.1101/2020.10.01.321638.

16. Brierley SM, Castro J, Harrington AM, Hughes PA, Page AJ, Rychkov GY, et al. TRPA1 contributes to specific mechanically activated currents and sensory neuron mechanical hypersensitivity. J Physiology 2011;589:3575–3593. doi:10.1113/jphysiol.2011.206789.

17. Conklin DJ, Guo Y, Nystoriak MA, Jagatheesan G, Obal D, Kilfoil PJ, et al. TRPA1 channel contributes to myocardial ischemia-reperfusion injury. Am J Physiol-Heart C 2019;316:H889–H899. doi:10.1152/ajpheart.00106.2018.

18. Zurborg S, Yurgionas B, Jira JA, Caspani O, Heppenstall PA. Direct activation of the ion channel TRPA1 by Ca2+. Nat Neurosci 2007;10:277–279. doi:10.1038/nn1843.

19. Andersson DA, Gentry C, Moss S, Bevan S. Transient receptor potential A1 is a sensory receptor for multiple products of oxidative stress. J Neurosci 2008;28:2485–2494. doi:10.1523/jneurosci.5369-07.2008.

20. Prosser BL, Ward CW, Lederer WJ. X-ROS Signaling: Rapid mechano-chemo transduction in heart. Science 2011;333:1440–1445. doi:10.1126/science.1202768.

21. Iribe G, Ward CW, Camelliti P, Bollensdorff C, Mason F, Burton RAB, et al. Axial stretch of rat single ventricular cardiomyocytes causes an acute and transient increase in Ca2+ spark rate. Circ Res 2009;104:787–795. doi:10.1161/circresaha.108.193334.

22. Cameron BA, Kai H, Kaihara K, Iribe G, Quinn TA. Ischemia enhances the acute stretch-induced increase in calcium spark rate in ventricular myocytes. Front Physiol 2020; 11:289. doi:10.3389/fphys.2020.00289.

23. Miura M, Wakayama Y, Endoh H, Nakano M, Sugai Y, Hirose M, et al. Spatial non-uniformity of excitation–contraction coupling can enhance arrhythmogenic-delayed afterdepolarizations in rat cardiac muscle. Cardiovasc Res 2008;80:55–61. doi:10.1093/cvr/cvn162.

24. Quinn TA, Granite S, Allessie MA, Antzelevitch C, Bollensdorff C, Bub G, et al. Minimum information about a cardiac electrophysiology experiment (MICEE): standardised reporting for model reproducibility, interoperability, and data sharing. Prog Biophysics Mol Biology 2011;107:4–10. doi:10.1016/j.pbiomolbio.2011.07.001.

25. Khokhlova A, Iribe G, Yamaguchi Y, Naruse K, Solovyova O. Effects of simulated ischemia on the transmural differences in the Frank–Starling relationship in isolated mouse ventricular cardiomyocytes. Prog Biophysics Mol Biology 2017;130:323–332. doi:10.1016/j.pbiomolbio.2017.05.011.

26. Zheng Q, Jockusch S, Zhou Z, Blanchard SC. The contribution of reactive oxygen species to the photobleaching of organic fluorophores. Photochem Photobiol 2014;90:448–454. doi:10.1111/php.12204.

27. Baumeister P, Quinn TA. Altered calcium handling and ventricular arrhythmias in acute ischemia. Clin Medicine Insights Cardiol 2016;10s1:CMC.S39706. doi:10.4137/cmc.s39706.

28. Keurs HEDJ ter, Boyden PA. Calcium and arrhythmogenesis. Physiol Rev 2007;87:457–506. doi:10.1152/physrev.00011.2006

29. Michailova A, Lorentz W, McCulloch A. Modeling transmural heterogeneity of KATP current in rabbit ventricular myocytes. Am J Physiol-Cell Ph 2007;293:C542–C557. doi:10.1152/ajpcell.00148.2006.

30. Calaghan SC, White E. The role of calcium in the response of cardiac muscle to stretch. Prog Biophysics Mol Biology 1999;71:59–90. doi:10.1016/s0079-6107(98)00037-6.

31. Keurs HEDJ ter, Wakayama Y, Miura M, Shinozaki T, Stuyvers BD, Boyden PA, et al. Arrhythmogenic Ca2+ release from cardiac myofilaments. Prog Biophysics Mol Biology 2006;90:151–171. doi:10.1016/j.pbiomolbio.2005.07.002.

32. Wang YY, Chang RB, Waters HN, McKemy DD, Liman ER. The nociceptor ion channel TRPA1 is potentiated and inactivated by permeating calcium ions. J Biological Chem 2008;283:32691–32703. doi:10.1074/jbc.m803568200.

33. Meents JE, Ciotu CI, Fischer MJM. TRPA1: a molecular view. J Neurophysiol 2019;121:427–443. doi:10.1152/jn.00524.2018.

34. Meents JE, Fischer MJM, McNaughton PA. Agonist-induced sensitisation of the irritant receptor ion channel TRPA1. J Physiology 2016;594:6643–6660. doi:10.1113/jp272237.

35. Wang Z, Xu Y, Wang M, Ye J, Liu J, Jiang H, et al. TRPA1 inhibition ameliorates pressure overload-induced cardiac hypertrophy and fibrosis in mice. Ebiomedicine 2018;36:54–62. doi:10.1016/j.ebiom.2018.08.022.

